# Zinc and laboratory emerge date impact honey bee gut microbiota and survival

**DOI:** 10.1101/2022.12.07.519529

**Authors:** Kilmer Oliveira Soares, Dylan Ricke, Celso José Bruno de Oliveira, Marcos Venâncio Lima, Ryan Mrofchak, Adriana Evangelista Rodrigues, Christopher Madden, Reed Johnson, Vanessa L. Hale

**Affiliations:** Department of Animal Science, College of Agricultural Sciences (CCA), Federal University of Paraiba (UFPB), Areia, PB, Brazil; Department of Entomology, Ohio Agricultural Research and Development Center, The Ohio State University, Wooster, Ohio, USA; Global One Health Initiative (GOHi), Ohio State University, Columbus, OH, 43210 USA; Department of Animal Science, College of Agricultural Sciences (CCA), Federal University of Ceara (UFC), Fortaleza, CE, Brazil; Department of Veterinary Preventive Medicine, College of Veterinary Medicine, Ohio State University (OSU), Columbus, OH, 43210 USA

**Keywords:** Honey bee, Gut Microbiota, Zinc, Emerge Day, Survival, Agrochemical

## Abstract

Honey bees (*Apis mellifera*) may be exposed to a wide variety of chemicals in the environment, including pesticides, antibiotics, and metals. Zinc, for example, is commonly included in fertilizers, pesticides, and feed additives, and is found in agricultural runoff. Honey bees can be exposed to zinc directly or indirectly by consuming zinc-contaminated nectar and pollen. However, there is a paucity of studies addressing the putative effects of zinc on honey bee’s health. In this study, we tested the effects of zinc on honey bee survivorship and gut microbiota. To evaluate survivorship, we exposed bees to six concentrations of zinc (0, 50, 100, 250, 500, or 1000 mg/L) and assessed survival daily for 10 days. To evaluate effects of zinc on gut microbiota, we exposed bees to 5 or 100 mg/L zinc. Bees were sampled before (day 0) and after zinc exposure (days 3, 6, and 9). Abdominal contents underwent DNA extraction and 16S rRNA sequencing (V3-V4) on an Illumina MiSeq. Sequences were filtered and processed through QIIME2 and DADA2. Zinc treatment had minimal effects on bacterial DNA concentrations and absolute cell counts while emerge date (the date a bee emerged from the brood comb) had a significant effect with decreased bacterial concentrations and cell counts observed at later emerge dates. Survival was only minimally impacted (>89% survival) at zinc concentrations up to 100 mg/L. Zinc had limited effects on overall gut microbial composition, diversity, and taxonomic abundances, with the greatest differences noted in the bee group exposed to the higher concentration of zinc (100 mg/L). In this group, several beneficial taxa (*Lactobacillus*, Rhizobiaceae, *Gilliamella*) were found at reduced abundances, while *Paenibacillus*, a potentially pathogenic taxa, was found at increased abundances. This suggests that zinc exposure, even at relatively low levels, may negatively impact honey bee health, even if survivorship is not dramatically impacted. Notably, emerge date effects were also observed in microbial composition. These results demonstrate the need to include assessments of honey bee gut microbiota in addition to other metrics of honey bee health and survivorship when evaluating the potential effects of agrochemicals on honey bees

## Introduction

Honey bees (*Apis mellifera*) are regularly exposed to a variety of chemicals in the environment, including pesticides, antibiotics, air pollutants, plastic additives, and metals (Roszko et al., 2016; Solayman et al., 2016; Soares et al., 2021). Exposure to substances like metals can occur via direct contact with airborne metals (Costa et al., 2019) or by consuming nectar and pollen contaminated with metals (Leita et al., 1996; Quinn et al., 2011). Although honey bee hives have long served as biomonitors of metal pollution (Bromenshenk et al., 1985; Leita et al., 1996), the effects of metals on honey bee health is relatively understudied. This is a major area of concern because metals, like zinc, are commonly found in fertilizers and foliar sprays, and can bioaccumulate, leading to toxicity in honey bees (Mortvedt and Gilkes, 1993; Montalvo et al., 2016; Hesketh et al., 2016; Hladun et al., 2016). Honey bees are economically important as food-producing animals and as pollinators across the planet (Kevan and Viana, 2003; Michener, 2007; Gisder and Genersch, 2017; Hung et al., 2018).

In recent years, a limited number of laboratory studies have shown that metals can negatively impact honey bee health including lifespan (Di et al., 2016, 2020; Hesketh et al., 2016), cognitive ability (Hladun et al., 2012; Søvik et al., 2015; Burden et al., 2016a, 2016b, 2019; Monchanin et al., 2021), and brood development (Hladun et al., 2013, 2016; Di et al., 2016, 2020). Metals also alter honey bee expression of genes involved in stress response and metal detoxification (Nikolić et al., 2016, 2019; Purać et al., 2019). Moreover, metal exposure results in more dead pupae within capped cells, lower worker weights, and increased metal accumulation in body tissue (Bromenshenk et al., 1991; Hladun et al., 2016). While certain metals (ex. cadmium and lead) have no known role within the body, others (ex. magnesium and zinc) are essential micronutrients that play key physiological processes in bees. However, excess exposure to essential metals, like zinc, can also result in toxic effects (Milivojević et al., 2015).

Zinc is frequently included in fertilizers (Mortvedt and Gilkes, 1993; Montalvo et al., 2016), and has been proposed as an active ingredient for plant antimicrobials (Naranjo et al., 2020). Zinc oxide nanoparticles have received attention as biosafe options for agricultural applications (Rajput et al., 2018; Mostafa et al., 2019). Additionally, zinc oxide is incorporated as a growth promoter in animal feed (Moynahan, 1979). Excess zinc is then excreted in animal waste, which can lead to zinc soil contamination (Nollet et al., 2007). Zinc can also bioaccumulate from soil into plants (Balafrej et al., 2020) and potentially into higher trophic levels, including florivores such as honey bees (Xun et al., 2017; Butt et al., 2018). Humans also excrete zinc (3-19 mg of zinc per day in feces, and 0.6-1.6 mg in urine; Drinker et al., 1927), which can make its way into agricultural fields via recycled wastewater (Gupta et al., 2019). Notably, zinc has been employed heavily as a supplement for the treatment and prevention of COVID-19, which could potentially translate into higher zinc levels in wastewater (Gordon and Hardigan, 2021; Michos and Cainzos-Achirica, 2021).

Honey bees have a wide foraging range (up to 10 km^2^; Beekman and Ratnieks, 2000) and may be exposed to zinc in fertilizers, pesticides, soils, plants, agricultural runoff, or recycled wastewater during foraging. However, the effects of zinc on honey bees is largely unknown. Importantly, metal exposure may affect the gut microbiome of exposed hosts (Syromyatnikov et al., 2020), and microbiota play a critical role in bee health (Hamdi et al., 2011; Engel et al., 2016; Raymann and Moran, 2018; Dosch et al., 2021). Certain microbes influence pesticide tolerance (Wu et al., 2020), behavior and signaling molecules (Kešnerová et al., 2017; Zhang et al., 2022), and inhibit the growth of fungal pathogens (Engel et al., 2016; Miller et al., 2020). The effects of pesticides on honey bee gut microbiota have been investigated in multiple studies (Kakumanu et al., 2016; Diaz and Larsen, 2018; Motta et al., 2018; Raymann et al., 2018; Nogrado et al., 2019; Paris et al., 2020; Cuesta-Maté et al., 2021; Hotchkiss et al., 2022), but the effects of metals, which can have antimicrobial properties (Lemire et al., 2013; Li et al., 2021), are largely unexplored (but see Rothman et al., 2019b).

Considering the current knowledge available, we hypothesize that both survival and gut microbial communities are negatively impacted by zinc exposure. Moreover, these are dose-dependent effects, with longer exposure periods and at higher concentrations of zinc leading to more pronounced deleterious effects. In the present study, we experimentally evaluated the effects of zinc exposure on the survival and gut microbial community of adult honey bees.

## Methods

### Experiment design

The experiment was conducted at the Ohio State University Honey Bee Research Laboratory at the Ohio Agricultural Research and Development Center (OARDC) in Wooster, OH (USA) between July 2020-October 2020. All honey bee hives used in this study were managed according to standard beekeeping practices (Honey Bee Health Coalition, 2019). The experimental design is presented in **Figure 1**.

**Figure 1.**
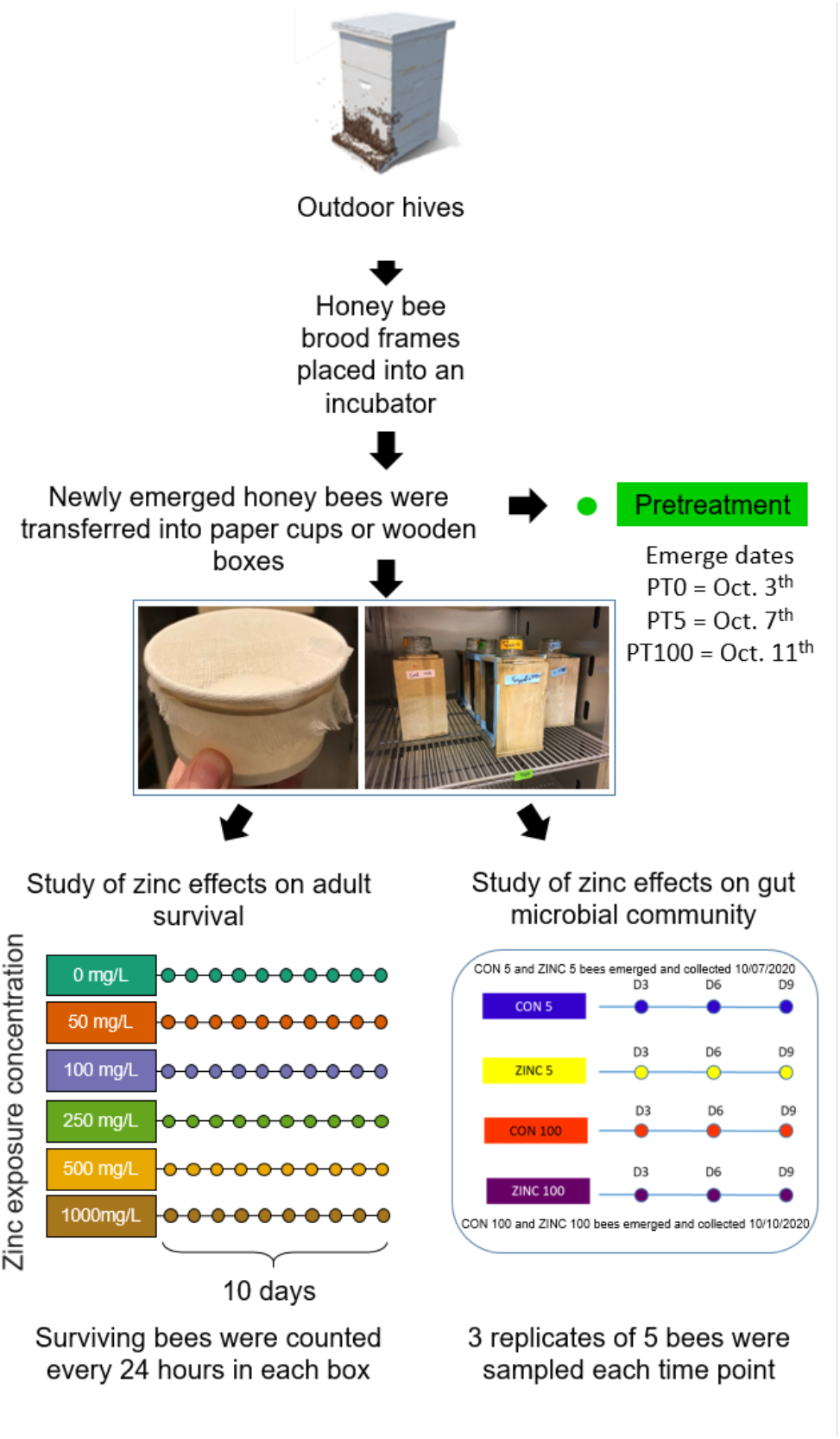
Experimental design. Honey bee brood frames were placed into a lab incubator. Newly emerged honey bees were collected daily. To evaluate the effects of zinc on the gut microbial community (right), bees were divided into wooden/mesh boxes and fed ad libitum with a sucrose-zinc solution at 5 mg/L (ZINC 5) or 100 mg/L (ZINC 100). Each treatment group was paired with a respective control group (CON 5 or CON 100) containing bees that emerged on the same date as the treatment group but that received no zinc (0 mg/L). Fifteen bees were collected from each box on Day 0 (Pretreatment - PT), Day 3 (D3), Day 6 (D6), and Day 9 (D9) for microbial analysis.

### Zinc oral toxicity assays

Zinc oral toxicity assays were conducted in July and August of 2020. A 1,000 mg/L stock solution of zinc was prepared by dissolving 112.4 mg of anhydrous zinc acetate (Sigma-Aldrich, St. Louis, MO) into 40 mL of 50% (w/w) sucrose solution. This stock solution was serially diluted with a 50% sucrose solution to yield experimental solutions containing 50, 100, 250, 500, or 1,000 mg/L of zinc (or 40.6, 81.3, 203.3, 406.5, and 813 mg/kg). A control solution with no added zinc was also prepared..

Assays followed standard protocols for assessing chronic oral toxicity in adult honey bees (OECD, 2017). Frames with emerging honey bees were collected from three healthy, queenright hives and stored in a frame box in a laboratory incubator (60-80% RH, 34°C). Newly emerged bees were collected from the box daily to ensure that all bees present on a given day were < 24 h old. For each trial, unanesthetized adult bees were brushed into a plastic bin and divided into groups of 17-23 bees (mean: 20.4, standard deviation: 1.2). All bees in each trial emerged from the brood frame on the same day. Groups were then transferred into 177 cm^3^ paper cups (Uniq Brand, Gilbert, AZ; **Figure 1**) and assigned to one of six treatment groups (0, 50, 100, 250, 500, or 1000 mg/L zinc). Bees were fed ad libitum from 1.5 mL microcentrifuge tubes that were modified to serve as feeders. Tubes were checked daily and refilled as needed. Surviving bees were counted every 24 hours (+/- 1.5 hours) for 10 days. Five replicates were performed for each treatment group. Replicates were omitted if their respective control groups (0 mg/L zinc) had ≤ 85% survival by the end of the 10-day trial (OECD, 2017). This resulted in each treatment group containing bees from 1-3 hives and 3-5 replicates (**Supplementary Table 1**).

### Analysis of oral toxicity data

To select an appropriate dose-response model, the US Environmental Protection Agency’s Benchmark Dose Software (Davis et al., 2011; USEPA, 2020) was used. This software recommended the probit model, which had the lowest Akaike Information Criterion among 12 competing models. Subsequently, a probit model of survival at day 10 over each concentration was generated with the *glm* function in the R package ‘stats’ (R Core Team, 2021). Additionally, models of survival over time were generated for each treatment group. These models were used to estimate the duration of exposure resulting in 50% mortality for each concentration (“lethal effect times,” or LT50s) with the *LC_probit* function in the R package ‘ecotox’ (Hlina et al., 2021).

To compare rates of survival between treatment groups, Kaplan-Meier survival estimates for each treatment group were calculated with the *survfit* function in the R package ‘survival’ (Therneau, 2021). Rates of survival between treatment groups were compared using the *pairwise_survdiff* function with a Bonferroni correction for multiple comparisons. This function performs multiple pairwise log-rank tests between the survival estimates of each treatment group.

### Zinc effects on gut microbiota

Gut microbiota experiments were performed in October of 2020. Honey bee brood frames were taken from a single outdoor hive and placed into an incubator with controlled temperature and humidity (60-80% RH, 34°C). On each day of the experiment, newly emerged honey bees were transferred into wooden/mesh boxes (∼1650 cm^3^) and provisioned with inverted glass feeding jars containing a 50% (w/w) sucrose solution with zinc at 5 mg/L (ZINC 5) or 100 mg/L (ZINC 100) (**Figure 1**). Boxes and jars were cleaned prior to this study by soaking into 30% bleach and rinsing thoroughly. The selected zinc concentrations (5 and 100 mg/L, or 4.07 and 81.3 mg/kg) were chosen to span the range of zinc concentrations previously reported in honey (Solayman et al., 2016). Notably, higher zinc concentrations (325 and 592 mg/kg) have been measured in bee propolis and pollen provisions, respectively (Leita et al., 1996; Moroń et al., 2012). Each zinc treatment group was paired with a respective control group (CON 5 or CON 100) containing bees that emerged on the same date as the treatment group but that received no zinc (0 mg/L). Fifteen bees were collected from each box on Day 0 (Pretreatment - PT), Day 3 (D3), Day 6 (D6), and Day 9 (D9) for gut microbial analysis. Bees were briefly anaesthetized at - 20°C before being individually transferred into sterile 1.5 mL microcentrifuge tubes containing 70% ethanol for preservation. Fifteen bees were sampled from each treatment and control group at each timepoint. Samples were then transported to the Ohio State University College of Veterinary Medicine in Columbus, Ohio for processing and gut microbial community analysis.

### DNA extraction, library preparation, and sequencing

Whole abdominal contents of five bees were pooled into a single tube for DNA extraction, which was performed using the Qiagen® PowerFecal® Pro DNA Isolation Kit, (Qiagen, Germany) following the manufacturer’s protocol with one alteration: A FastPrep-24 5G beat beater (MP Biomedicals, USA) with a setting of 6 m/s for 40s was used in place of the vortex adapter. The final elution step was performed in 50 μl to maximize DNA concentration. After extraction, DNA was assessed for concentration, purity, and integrity using the a fluorometer (Qubit 4, Thermo Fisher, USA) and a microvolume spectrophotometer (NanoDrop™, Thermo Fisher, USA).

Sequencing and library preparation (16S rRNA) was performed by Novogene Technology Co. Ltd. (Beijing, China). 16S rRNA amplicon library was constructed from each sample using primers 341F: 5’-CCTAYGGGRBGCASCAG-3′ and 806R: 5’-GGACTACNNGGGTATCTAAT-3′ targeting the V3-V4 region of the 16S rRNA gene (Jia et al., 2017). PCR reaction conditions were as follows: initial denaturation at 95°C for 3 min, followed by 25 cycles at 95°C for 30 s, 55°C for 30 s, and 72°C for 30s and a final extension at 72°C for 5 min. Barcodes were added to the primer sequences. The sequencing libraries were quantified by fluorometry (Qubit2.0, Life Invitrogen) and an Agilent Bioanalyzer 2100 system prior to sequencing. An Illumina HiSeq 2500 platform was employed to generate 2 × 250 bp paired-end reads. Negative controls including blanks from the extraction kit and samples from each of the feeds also underwent extraction and sequencing.

### Bacterial DNA quantification

Bacterial DNA concentrations were quantified through qPCR: Bacterial DNA was amplified using 16S rRNA universal bacterial primers and probes according to Nadkarni et al. (2002) using a QuantStudio 3 Real-Time PCR System (Applied Biosystems, Thermo Fisher Scientific, Carlsbad, CA, USA). For the reaction, 300 nM of forward primer (5’ – TCCTACGGGAGGCAGCAGT – 3’), 300 nM of reverse primer (5’ – GGACTACCAGGGTATCTAATCCTGTT – 3’), and 175 nM of probe ((6FAM) – 5’ – CGTATTACCGCGGCTGCTGGCAC – 3’ – (TAMRA)) were added. Cycling parameters were as follows: 50 °C for 2 min and 95 °C for 10 min followed by 40 cycles of 95 °C for 15 s and an annealing and extension step of 60 °C for 1 min (Nadkarni et al., 2002). qPCR was performed in triplicate and to be included in analyses, at least two replicates per sample had to amplify. Samples containing replicates with greater than 3% variation were removed from analysis. Following qPCR, cycle thresholds were log_10_-transformed using the following equation based on a standard curve generated using an *Escherichia coli* isolate (Mrofchak et al., 2021): y = -5.329x + 36.504 in which y is the cycle threshold and x is the log_10_-transformed DNA concentration. The antilog of each sample was then used to calculate the bacterial DNA concentration in each sample.

### Sequence Data Processing and Statistical analysis

The raw demultiplexed paired-end sequences were processed using QIIME 2-2020.2 (Bolyen et al., 2019). Paired-end reads were filtered, denoised, and truncated to a length of 248 base pairs, and then parsed for non-chimeric sequences using DADA2, producing Amplicon Sequence Variants (ASV) (Callahan et al., 2016). Sequences were aligned using “qiime fragment-insertion sepp” for phylogenetic analysis (Matsen et al., 2012). Taxonomy was assigned in QIIME2 using SILVA version 132, with a 99% similarity threshold using the full length 16S rRNA gene classifier (Quast et al., 2013; Yilmaz et al., 2014). Negative control samples were examined for potential contaminant taxa and reads were then subjected to in-silico decontamination using the Decontam R package version 1.12.0 (Davis et al., 2017). Microbial diversity (alpha diversity) was assessed using observed features (richness), Shannon (richness and abundance), Faith’s PD (phylogenetic diversity) and Pielou’s (evenness) diversity indices. ANOVAs were used to compare diversity between groups using R version 4.1.0 (Ripley, 2001). After testing for normality using a Shapiro-Wilk test, the means were compared using Tukey or Kruskal-Wallis tests at 5% probability.

Microbial composition (beta diversity) was evaluated using Bray-Curtis, Jaccard, and weighted and unweighted UniFrac distances in QIIME 2-2020.2. Microbial community similarity and dissimilarity were visualized with principle coordinate analysis (PCoA) plots using the Emperor plugin 2020.2.0 (Vázquez-Baeza et al., 2017). PERMANOVAs were employed as recommended (Anderson, 2001) to test for differences in microbial composition between experimental groups: (Pretreatment (PT), zinc at 5 mg/L (ZINC 5), zinc at 100 mg/L (ZINC 100), and their respective matched control groups that received 0 mg/L zinc (CON 5, CON 100). Microbial communities were also compared over time (Day 0 – Pretreatment (PT), and Days 3 (D3), 6 (D6), and 9 (D9)).

Taxa that were differentially abundant between the treatment groups were identified using an analysis of composition of microbiomes (ANCOM) (Mandal et al., 2015). We also used the “qiime feature-table core-features” command to identify core taxa present in 100% of the samples. The relative and absolute abundances of differentially abundant microbes were compared by treatment using one-way ANOVAs followed by Tukey or Kruskal-Wallis Rank Sum tests and pairwise comparisons using pairwise Wilcoxon Rank Sum Tests. Normality of data was assessed by the Shapiro-Wilk test. A *P*-value < 0.05 was used in the statistical tests for significance.

## Results

### Zinc effects on honey bee survival

Honey bee survival exhibited a clear dose-response relationship with zinc concentration (log-rank test, p < 0.0001, **χ**^2^ = 1073, df = 5, **Figure 2, Supplementary Figure 1**). At 0, 50, and 100 mg/L zinc, mean rates of survival were 99%, 91%, and 89%, respectively. Survival decreased steeply at 250 mg/L zinc exposure and was 0% at 1000 mg/L (**Supplementary Table 2**). Accordingly, LT50s (lethal effect times to reach 50% mortality) decreased with increasing zinc concentrations (**Supplementary Table 3**). Pairwise comparisons between treatment groups revealed statistically significant differences in overall survival between all groups (p << 0.05) except the 50 and 100 mg/L groups (p ∼ 1) (**Supplementary Table 4**). Similarly, the 95% confidence intervals of probit LT50 estimates only overlapped for the 50 and 100 mg/L groups (**Supplementary Table 3**). Based on these survivorship results, we chose to test the effects of two field-relevant concentrations of zinc (5 mg/L and 100 mg/L) on the honey bee gut microbiota. While zinc concentrations up to 100 mg/L had relatively small effects on survival, gut microbial community shifts could impact honey bee health in other ways.

**Figure 2.**
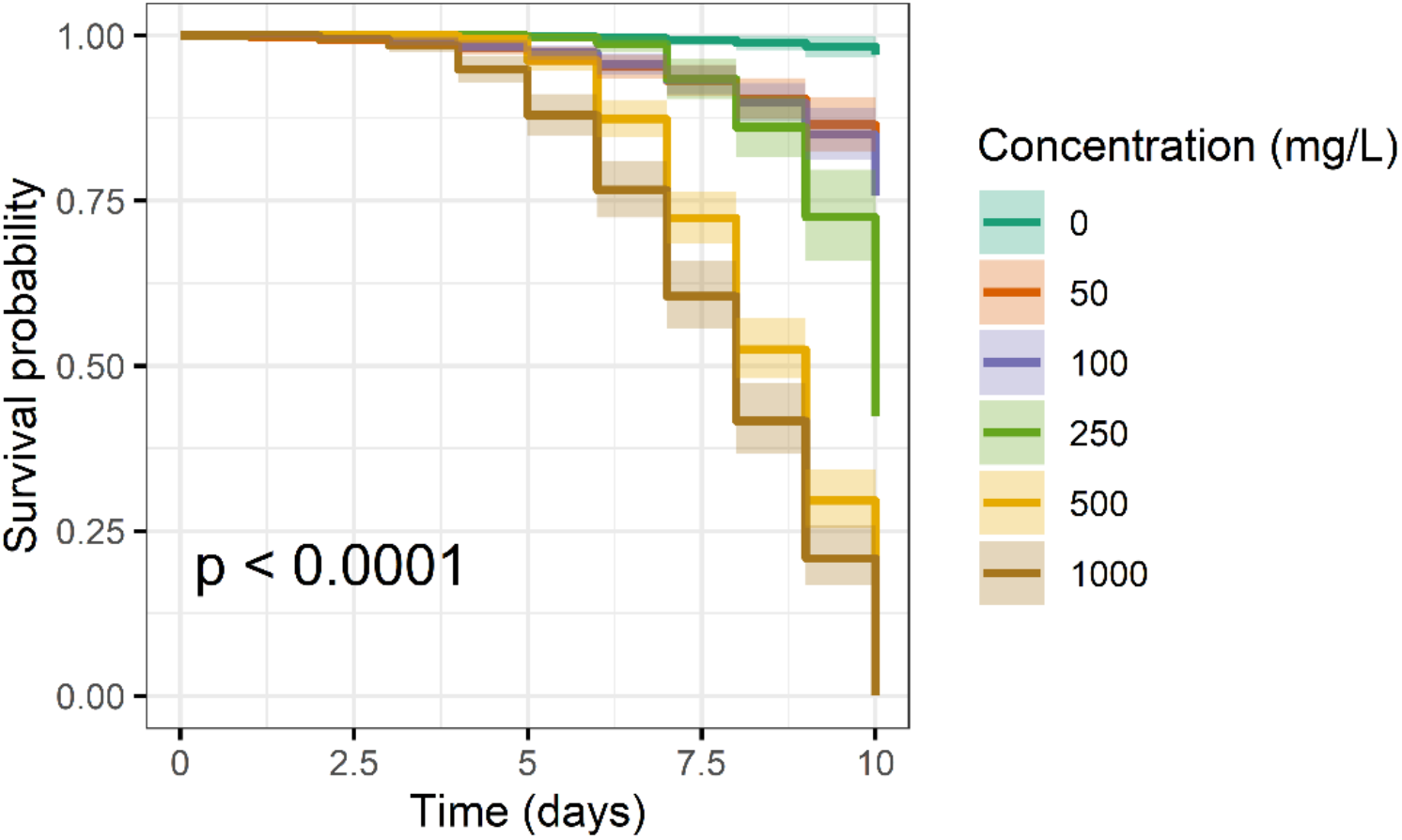
Kaplan-Meier curves of honey bee survival at varying zinc concentrations over 10 days. Zinc concentrations included 0 (negative control), 50, 100, 250, 500, and 1000 mg/L. Increased mortality was observed with increasing zinc concentrations. Lines and shaded regions represent median survival predictions and 95% confidence regions, respectively. Across all groups, zinc concentration had a statistically significant effect on survival (log-rank test, p < 0.0001, χ^2^ = 1073, df = 5). When focusing on just the field-relevant concentrations (0, 50, and 100 mg/L), concentration still had a statistically significant effect on survival (log-rank test, p < 0.0001, χ^2^ = 45.9, df = 2).

### Zinc effects on honey bee bacterial DNA concentration and absolute cell counts

To assess the effects of zinc on overall gut bacterial load, we quantified total bacterial DNA concentrations and absolute bacterial cell counts via qPCR. Five samples exhibited greater than 3% variation in cycle threshold values between all replicates and were removed from analyses. This included two ZINC 5 samples from day 6, one CON 5 sample from day 6 and day 9, and one ZINC 100 sample from the day 3. In cases where only one of the three replicates exceeded 3% variation in cycle threshold value, the replicate with > 3% variation was excluded from analysis, and the remaining two replicates were retained and averaged. Cycle threshold values were then converted to bacterial DNA concentrations based on an equation generated from an *Escherichia coli* standard curve in our laboratory (Mrofchak et al., 2021).

Quantifiable bacterial DNA concentrations ranged from 0.0004 ng/ul to 5.87 ng/ul (**Supplementary Table 5**). Concentrations and absolute cell counts differed significantly by treatment (PT, ZINC 5, ZINC 100, CON 5, CON 100) (Bacterial concentration: Kruskal Wallis = 14.7, df = 4, p = 0.005; absolute cell count: Kruskal Wallis = 14.7, df = 4, p = 0.005), but no pairwise comparisons were significant (Wilcoxon rank sum exact test, all p > 0.05; **Figure 3**). Interestingly, both CON 5 and ZINC 5 groups had higher concentrations compared to CON 100 and ZINC 100 groups. CON 5 and ZINC 5 bees all emerged on the same date (October 7, 2021), which was 4 days before all CON 100 and ZINC 100 bees, which emerged on October 11, 2021. This led us to wonder if gut bacterial DNA concentrations decreased over time under laboratory conditions, and if bees with later emerge days had lower overall microbial loads. To test this, we examined bees in the PT group.

**Figure 3.**
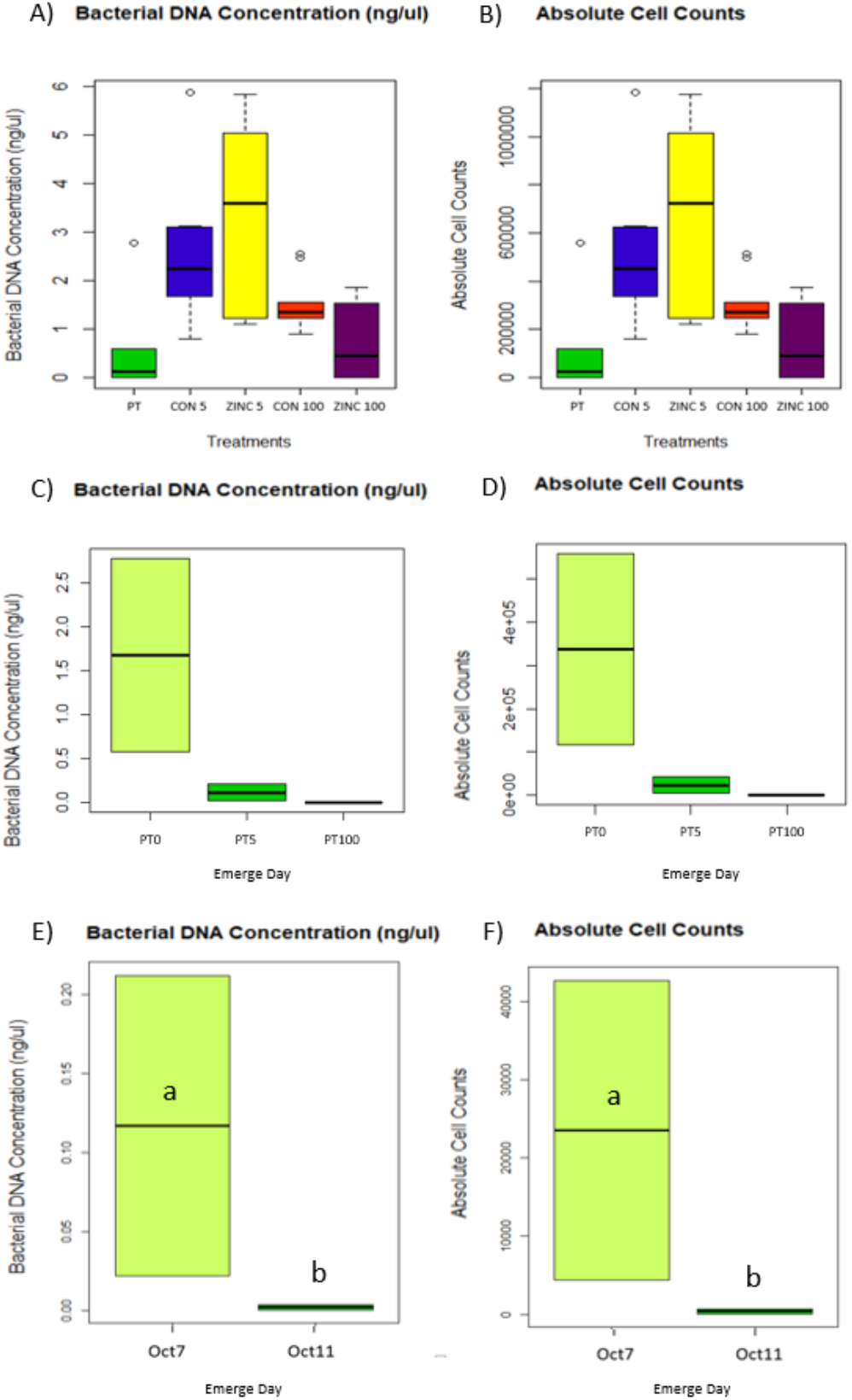
Bacterial DNA concentrations and absolute cell counts. **A**) Bacterial DNA concentrations and **B**) absolute cell counts differed significantly across treatment groups (Kruskal-Wallis, p = 0.005). However, no pairwise comparisons were significant (Wilcoxon Rank Sums, p > 0.05). **C**) Pretreatment (PT) bacterial DNA concentrations and **D**) absolute cell counts showed a non-significant decrease over time by emerge day (Bacterial DNA concentration: Kruskal-Wallis chi-squared = 4.57, df = 2, p-value = 0.102; absolute cell count: Kruskal-Wallis chi-squared = 4.57, df = 2, p-value = 0.102). When we combined all bees that emerged on the same day across all treatments and times, we found that **E**) bacterial DNA concentrations and **F**) absolute cell counts were significantly greater in bees that emerged on October 7^th^ as compared to October 11th (Bacterial DNA concentrations Kruskal-Wallis chi-squared = 6.15, df = 1, p-value = 0.013; absolute cell counts Kruskal-Wallis chi-squared = 6.1501, df = 1, p-value = 0.013); Box plot shows outliers, first and third quartiles (lower and upper edges), and highest, lowest and median values (horizontal black dash). PT = All Pretreatment (Day 0) bees including those that emerged on Oct. 3, 7, and 11; PT0 = Pretreatment bees that emerged on Oct. 3; PT5 = Pretreatment bees that emerged on Oct 7^th^ – the same day as CON 5 and ZINC 5 bees; PT100 = Pretreatment bees that emerged on Oct. 11^th.^ – the same day as CON 100 and ZINC 100 bees); CON 5 = Bees that emerged on the same day as ZINC 5 bees but treated with no zinc; ZINC 5 = Bees treated with 5 mg/L zinc; CON 100 = Bees that emerged on the same day as ZINC 100 bees but treated with no zinc; ZINC 100 = Bees treated with 100 mg/L zinc.

When emerge dates of PT bees were considered (PT 0 = bees that emerged on Oct. 3; PT 5 = bees that emerged on Oct. 7^th^, the same date as CON 5 and ZINC 5 bees; PT 100 = bees that emerged on Oct. 11^th^, the same date as CON 100 and ZINC 100 bees), we observed a clear decline in DNA concentrations and absolute cell counts over time by emerge date; although, these differences were still not significant (Bacterial DNA concentration: Kruskal-Wallis chi-squared = 4.5714, df = 2, p-value = 0.1017; absolute cell count: Kruskal-Wallis chi-squared = 4.57, df = 2, p-value = 0.102; **Figure 3c,d**). We then averaged data from all bees that emerged on Oct. 7th (PT5, CON 5, ZINC 5), and all bees that emerged on Oct. 11^th^ (PT100, CON 100, ZINC 100) and, in that case, we observed a significant difference in bacterial DNA concentrations and absolute cell counts between emerge day (Bacterial DNA concentration: Kruskal-Wallis chi-squared = 6.15, df = 1, p-value = 0.013; Absolute cell count: Kruskal-Wallis chi-squared = 6.15, df = 1, p-value = 0.013; **Figure 3e,f**).

### 16S rRNA sequencing

We obtained a total of 4,854,562 raw reads across all samples. Samples averaged 112,896 reads per sample and ranged from 67,922 to 148,812 reads. After the denoising process, 4,165,574 (85.86%) reads were retained for downstream analyses. Five putative contaminant taxa (*Romboutsia* sp. CE17, *Methylobacterium* sp., *Cutibacterium acnes* strain 3265, *Methylorubrum extorquens* strain B44 and Uncultured bacterium clone A1435) were identified and removed using the decontam R package (Ripley, 2001) (**Supplementary Table 6**). Reads identified as chloroplasts, mitochondria, unassigned, and eukaryota were also removed from all samples. In total, reads were classified into 1,746 amplicon sequence variants (ASVs) which aligned to 413 different taxa. Samples were rarefied at 3,000 reads.

### Zinc effects on honey bee gut microbial composition and diversity

Bee gut microbial composition was significantly different across treatment groups (PT, ZINC 5, ZINC 100, CON 5, CON 100) but not by time (D0/PT, D3, D6, D9) (PERMANOVA: Treatment - Bray Curtis R^2^= 0.280, *p =* 0.001; Jaccard Index: R^2^= 0.198, *p =* 0.001; Time - Bray Curtis R^2^= 0.026, *p =* 0.202; Jaccard Index: R^2^= 0.020, *p =* 0.202; **Figure 4, Supplementary Figure 2, Supplementary Tables 7-10**). The Bray Curtis distance metrics revealed that CON 100 and ZINC 100 groups did not differ significantly from each other in terms of microbial composition, and neither did CON 5 or ZINC 5 groups. However, both CON 100 and ZINC 100 groups differed significantly from CON 5 and ZINC 5 groups (**Supplementary Table 9**). These results again suggested an “emerge day” effect and indicated that emerge day and not zinc was driving the main differences observed between groups. We then analyzed microbial composition by emerge day (Oct. 7 vs. Oct. 11) and found significant differences between groups across all indices (PERMANOVA: Bray Curtis R^2^= 0.17, *p =* 0.001; Jaccard R^2^= 0.08, *p =* 0.001; Unweighted Unifrac R^2^= 0.08, *p =* 0.00; Weighted Unifrac R^2^= 0.17, *p =* 0.001).

**Figure 4.**
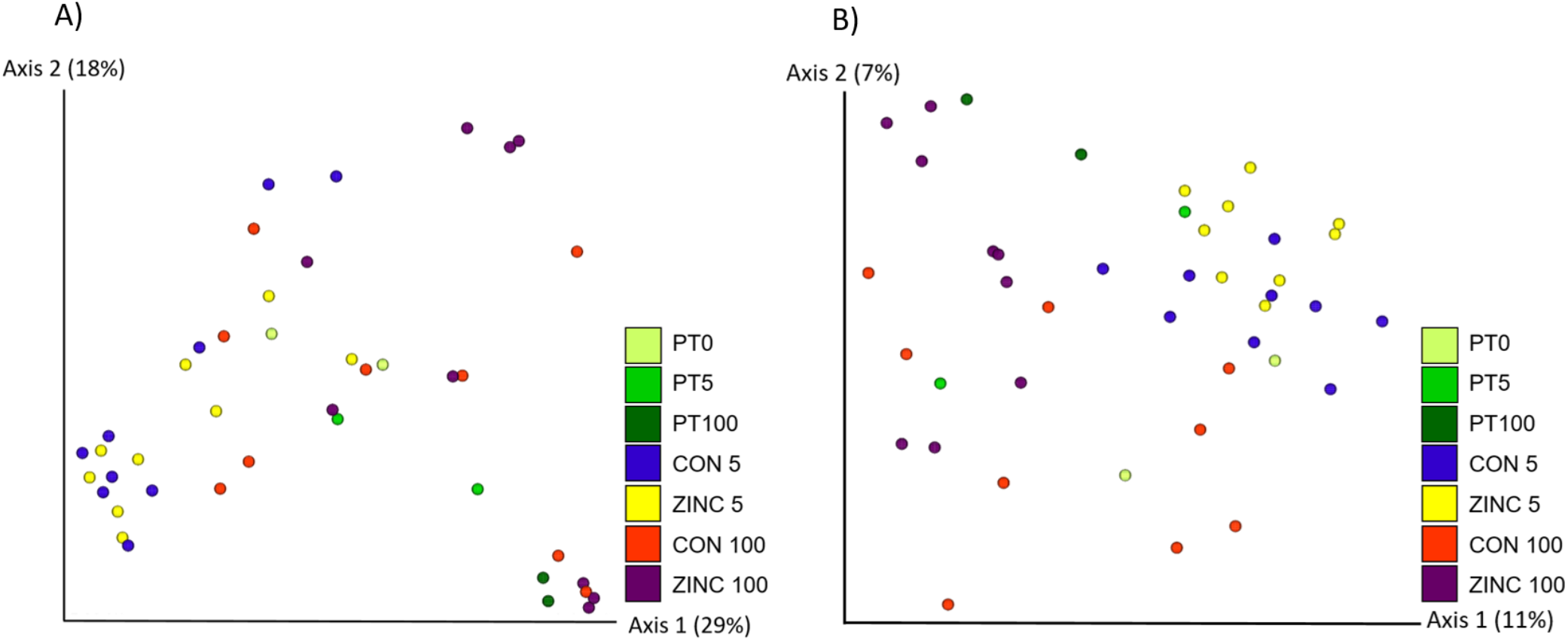
Bee gut microbial composition by treatment. Bee gut microbial composition differed significantly by treatment based on **A**) Bray Curtis (PERMANOVA: *p =* 0.001) and **B**) Jaccard distance matrices (PERMANOVA: *p =* 0.001). These differences were largely driven by emerge day as opposed to zinc treatment (also see **Supplementary Tables 9,10**). PT0 = Pretreatment bees that emerged on Oct. 3; PT5 = Pretreatment bees that emerged on Oct 7^th^ – the same day as CON 5 and ZINC 5 bees; PT100 = Pretreatment bees that emerged on Oct. 11^th.^ – the same day as CON 100 and ZINC 100 bees); CON 5 = Bees that emerged on the same day as ZINC 5 bees but treated with no zinc; ZINC 5 = Bees treated with 5 mg/L zinc; CON 100 = Bees that emerged on the same day as ZINC 100 bees but treated with no zinc; ZINC 100 = Bees treated with 100 mg/L zinc.

To compare microbial diversity across groups, we used Shannon and Pielou’s Indices after first testing data for normality using the Shapiro Wilk Test. Microbial diversity did not differ significantly by treatment groups or by time (Two way ANOVA: Treatment - Shannon Index *p =* 0.183; Pielou’s Index *p =* 0.226; Time (D0, D3, D6, D9) - Shannon Index *p =* 0.105; Pielou’s Index *p =* 0.08; **Figure 5a,b**; **Supplementary Figure 3a,b)**. To test for emerge day effects, we also compared microbial diversity of PT bees by emerge date and found no significant differences (Emerge Day - Shannon Index *p =* 0.671; Pielou’s Index *p =* 0.952 **Figure 5c,d; Supplementary Figure 3c,d**). Further, we compared microbial diversity of all bees that emerged on Oct. 7 (PT5, CON 5, ZINC 5) to all bees that emerged on Oct. 11 (PT100, CON 100, ZINC 100) and again found no significant differences (ANOVA: Treatment - Shannon Index *p =* 0.216; Pielou’s Index *p =* 0.376; **Figure 5e,f**).

**Figure 5.**
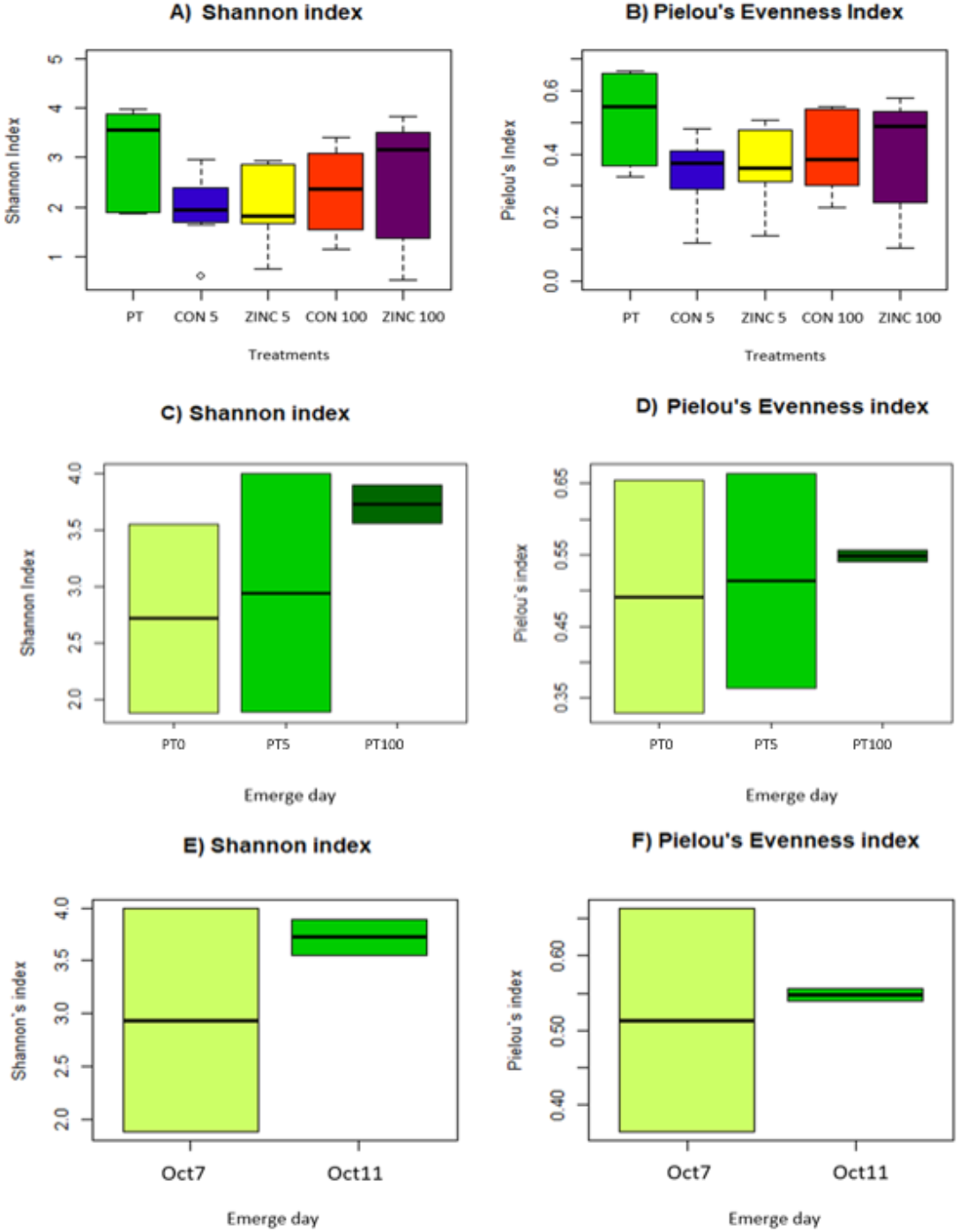
Microbial diversity by treatment. Gut microbial diversity and evenness did not differ significantly by treatment (**A,B**) or by emerge day (**C,D**) as measured by the Shannon (**A,C**) or Pielou’s evenness index (**B,D**) (ANOVA: Treatment - Shannon Index *p =* 0.216; Pielou’s Index *p =* 0.211; Emerge Day - Shannon Index *p =* 0.671; Pielou’s Index *p =* 0.952. When we combined all bees that emerged on the same day across all treatments and times, we still found no significant difference in microbial diversity by emerge day (**E,F**) (ANOVA: Emerge day - Shannon Index *p =* 0.216; Pielou’s Index *p =* 0.376). Box plots show outliers, first and third quartiles (lower and upper edges), and highest, lowest and median values (horizontal black dash). PT = All Pretreatment (Day 0) bees including those that emerged on Oct. 3, 7, and 11; PT0 = Pretreatment bees that emerged on Oct. 3; PT 5 = Pretreatment bees that emerged on Oct 7^th^ – the same day as CON 5 and ZINC 5 bees; PT 100 = Pretreatment bees that emerged on Oct. 11^th.^ – the same day as CON 100 and ZINC 100 bees); CON 5 = Bees that emerged on the same day as ZINC 5 bees but treated with no zinc; ZINC 5 = Bees treated with 5 mg/L zinc; CON 100 = Bees that emerged on the same day as ZINC 100 bees but treated with no zinc; ZINC 100 = Bees treated with 100 mg/L zinc.

### Core microbiota and differentially abundant taxa

A core microbiota analysis was performed to identify taxa present in 100% of the samples across all treatments and times. Four core taxa were identified including: *Lactobacillus, Bifidobacterium, Commensalibacter* and a taxon in the Rhizobiaceae family (**Supplementary Table 11**). These taxa accounted for 8% of all taxa in the dataset.

Six taxa (at the genera level) were identified as differentially abundant between treatment groups by ANCOM: *Lactobacillus, Gilliamella, Paenibacillus*, and three taxa from the families Rhizobiaceae, Enterobacteriaceae and Microbacteriaceae (**Figure 6**; **Supplementary Figure 4**; **Supplementary Table 12**). Significantly decreased abundances of Enterobacteriaceae and increased abundances in *Paenibacillus* and Microbacteriaceae were observed in the ZINC 100 group compared to the CON 100 group (Kruskal-Wallis test on absolute abundances: Enterobacteriaceae p = 0.0003; Microbacteriaceae p = 0.0001; **Figure 6, c,e,f**; Relative abundances: Enterobacteriaceae p = 0.00007; *Paenibacillus* p = 0.0005; Microbacteriaceae p = 0.0007; **Supplementary Figure 4, c,e,f**). For *Lactobacillus*, the Rhizobiaceae taxa, and *Gilliamella*, there were slight but non-significant decreases in relative and absolute abundances of these taxa in the zinc treated groups (ZINC 5 and ZINC 100). Notably, all CON 5 and ZINC 5 bees (emerge date: Oct. 7) contained greater relative abundances of *Lactobacillus*, the Rhizobiaceae taxa, and *Gilliamella* as compared to all CON 100 and ZINC 100 bees (emerge date Oct 11), suggesting unique emerge day impacts on some microbial taxa distinct from the effects of zinc.

**Figure 6.**
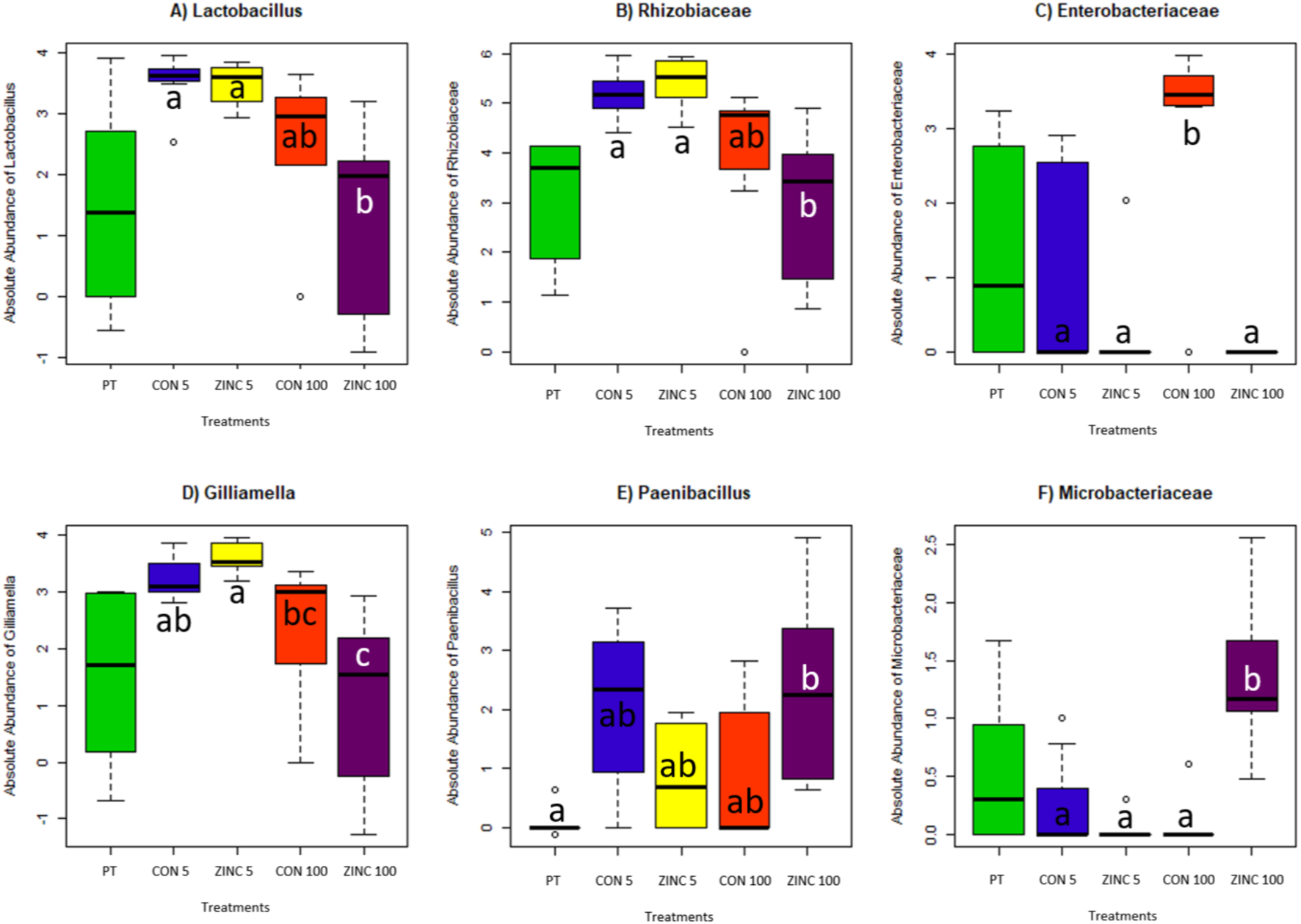
Differentially abundant microbiota by treatment. Absolute abundances of bacteria that were differentially abundant (ANCOM) at the L7 (roughly species) level between groups. Each species is listed at the lowest taxonomic level that could be identified based on reference sequences: **A**) *Lactobacillus*, **B**) *Bifidobacterium*, **C**) Rhizobiaceae, **D**) *Gilliamella*, **E**) *Tyzzerella*, **F**) *Streptomyces*, **G**) *Paenibacillus*, **H**) Enterobacteriaceae, **I**) Proteobacteria. Box plot shows outliers, first and third quartiles (lower and upper edges), and highest, lowest and median values (horizontal black dash). Groups that share the same letter do not differ significantly (Wilcoxon rank sum test with continuity correction). PT = All Pretreatment (Day 0) bees including those that emerged on Oct. 3, 7, and 11; CON 5 = Bees that emerged on the same day as ZINC 5 bees but treated with no zinc; ZINC 5 = Bees treated with 5 mg/L zinc; CON 100 = Bees that emerged on the same day as ZINC 100 bees but treated with no zinc; ZINC 100 = Bees treated with 100 mg/L zinc.

## Discussion

Our results demonstrate that low level zinc exposure had limited impacts on honey bee survival and overall gut microbial composition and diversity over time. However, we observed significant shifts in a few core and non-core bacterial abundances by treatment and emerge day. These zinc-induced gut microbial changes have potentially negative implications for honey bee nutrient metabolism and pathogen resistance. Importantly, the gut microbial community effects observed by emerge day indicate that emerge day must be accounted for as a variable in laboratory-based experiments involving honey bees.

### Zinc and honey bee survival

Each concentration of zinc tested in the oral toxicity assays produced a significant, negative effect on honey bee survival. Notably, this included zinc at concentrations of 50 and 100 mg/L, which are within the range of zinc concentrations reported from honey (Solayman et al., 2016). Zhang et al. (2015) reported similarly decreased survival of adult worker bees when fed zinc at concentrations of 45 mg/kg. Interestingly, they also observed increased longevity when honey bees were fed zinc at lower concentrations (30 mg/kg), which also resulted in increased antioxidant activity of the enzyme Cu-Zn superoxide dismutase (Cu-Zn SOD). Taken together, this suggests that honey bees could benefit from added zinc when zinc concentrations are low (i.e. ≤ 30 mg/kg), but zinc can have detrimental effects on adult honey bees at higher concentrations. Notably, while organic zinc acetate was used in this study, different formulations of zinc (e.g. zinc-methionine, zinc sulfate, zinc oxide, zinc nanoparticles) can yield different and highly variable results (De Barros et al. 2021). For example, a previous study reported a 20% reduction in survival and a 30% reduction in brain weight of honey bees exposed to just 0.8 mg/mL zinc oxide for ten days, indicating that zinc oxide may be significantly more toxic than zinc acetate (Milivojević et al. 2015).

### Zinc, emerge date, and honey bee gut microbiota

Zinc treatment (5 mg/L or 100 mg/L) had minimal effects on honey bee gut bacterial DNA concentrations and absolute cell counts while emerge date had a more pronounced, although still non-significant, effect with decreased bacterial concentrations and cell counts at later emerge dates. In a previous study, honey bees reared with decreased bacterial loads (<10^5^) demonstrated decreased weight gain, altered sensitivity to sugar, and significant changes in gut pH, oxygen gradient, and metabolite profiles (Zheng et al., 2017). This suggests that laboratory emerge day could have critical effects on honey bee health distinct from any experimental treatments being tested in the lab.

Zinc had minimal effects on overall gut microbial composition and diversity (**Figure 4, 5**). Although non-significant, a slightly larger shift in microbial composition was observed in the higher zinc exposure group (ZINC 100) compared to its control (CON 100) while essentially no shift was observed in the ZINC 5 group compared to its control (CON 5). Emerge day, on the other hand, had significant effects on microbial composition (**Supplementary Tables 7, 8, 9 and 10**).

Among the differentially abundant taxa identified between treatment groups, Enterobacteriaceae was found at high abundances in the CON 100 group as compared to ZINC 100 group and compared to the CON 5, ZINC 5, and PT groups. Enterobacteriaceae have been linked with dysbiosis and increased mortality in honey bees, and negatively correlated with Lactobacillaceae, which are considered beneficial bacteria that stimulate the honey bee innate immune system (Budge et al., 2016; Kwong and Moran, 2016; Bleau et al., 2020). One possibility to explain the high abundances of Enterobacteriaceae in the CON 100 but not ZINC 100 group is that Enterobacteriaceae expansion is supported at later emerge days but zinc exposure limited Enterobacteriaceae growth in the ZINC 100 group. Newly emerged honey bees consume probiotic-rich bee bread in the hive, which is produced through the bacterial fermentation of pollen. Anaerobic bacteria within the Enterobacteriaceae family play a key role in pollen fermentation to bee bread (Mattila et al., 2012). It is possible that the microbial populations present within bee bread - including Enterobacteriacea - change over time in the laboratory due to differing environmental conditions from the hive or a lack of regular contact with nurse bees. These changes in bee bread may then directly or indirectly facilitate an opportunistic bloom of Enterobacteriacea in the honey bee gut.

Although the ZINC 100 group did not show increased abundances of Enterobacteriacea, we did observe significant shifts in other microbial abundances including an unidentified *Paenibacillus* species that was increased in the ZINC 100 group as compared the CON 100 group. Some members of the *Paenibacillus* genus, such as *P. larvae*, the causative agent of American foulbrood (AFB), are considered honey bee pathogens (Grady et al., 2016). *P. larvae* spores are commonly found in hives, and are vectored by adult honey bees that are resistant to infection but can transmit the spores to new brood (Riessberger-Gallé et al., 2001). Exposure to high concentrations of zinc may allow *Paenibacillus* species like *P. larvae* to proliferate in the honey bee gut, which could pose increased risks for infection and transmission of diseases like AFB.

A Microbacteriaceae species was also found at significantly increased abundances in the ZINC 100 group as compared to the CON 100 group and the PT, CON 5, and ZINC 5 groups. Microbacteriaceae belong to the Actinobacteria family which is generally found at low relative abundances in the bee gut (Keller et al., 2013; Rothman et al., 2019a; Kačániová et al., 2020; Prado et al., 2022). Interestingly, one Microbacteriaceae species, isolated from a willow tree growing in soil heavily contaminated with zinc and cadmium, demonstrated heavy metal resistance (Corretto et al., 2017). This suggests that bacteria within this family may be uniquely suited to survive or even grow at high zinc concentrations. Future efforts involving isolation and genome sequencing of the honey bee Microbacteriacea are required to confirm this.

Decreased abundances of *Lactobacillus*, Rhizobiaceae, and *Gilliamella* were observed both in ZINC 100 bees as compared to CON 100 bees and in the Oct. 11 emerge day bees as compared to the Oct. 7 emerge day bees. These differences were not significant but hint at potential effects of zinc and emerge day on these taxa, which play important roles in bee and hive health. *Lactobacillus* bacteria convert pollen into nutrients more accessible to the bee host (Kešnerová et al., 2017). Rhizobiaceae taxa are consistently found at greater abundances in healthy bees relative to bees from collapsing colonies; although, their function has yet to be fully defined (Cornman et al., 2012). Finally, *Gillamella* species produce a biofilm on the gut wall that aids in pathogen defense (Engel et al., 2012).

In this study, zinc had limited effects on overall gut microbial composition, diversity, and taxonomic abundances, with the greatest differences noted particularly in the groups exposed to the higher concentration of zinc (100 mg/L). Effects linked to zinc included decreased abundances of several taxa including Enterobacteriacea (significant), and *Lactobacillus*, Rhizobiaceae, and *Gilliamella* (all non-significant). Zinc has antibacterial properties that may have driven some of these decreases (Almoudi et al., 2018). However, significant increases in other taxa, including *Paenibacillus* and Microbacteriacea, were also observed in the ZINC 100 group. The *Paenibacillus* taxa could represent a potential pathogen bloom resulting from altered microbiota, while the Microbacteriacea enrichment could be associated with zinc resistance in this taxon. Bees exposed to 100 mg/L zinc also demonstrated increased mortality, indicating that overall, zinc exposure at this field-relevant concentration negatively impacted bee health. These negative effects could be the result of zinc toxicity directly on host (bee) cells, or they may be mediated through changes in gut microbiota that alter other aspects of health such as immune dynamics and pathogen resistance (Budge et al., 2016; Powell et al., 2016; Kwong et al., 2017; Bleau et al., 2020; Miller et al., 2020). Importantly, zinc, in various formulations (Kolodziejczak-Radzimska and Jesionowski, 2014; Naranjo et al., 2020), continues to be incorporated into bactericides designed for application on plants. These products could have serious negative impacts on honey bees, and continued evaluation of safe dosing ranges and formulations should include honey bee gut microbial analyses.

Independent of zinc, emerge day clearly affects gut microbial composition, microbial load, and, to a lesser degree, in taxonomic abundances. Honey bees that emerged on different days were likely exposed to differing pre- and probiotic elements on the surface of the brood frame due to aging of the wax, changes over time in colony residues (bee bread, feces, etc.), lack of new bacteria typically introduced by other bees moving within the hive, or the removal of residues by previously-emerged bees. This was an unexpected finding, and it highlights a clear need to control for emerge day in laboratory experiments like this one. Additional studies are needed to determine the type and source of emerge day effects, as well as the potential health impacts of these effects. Determining if emerge day also impacts microbiota under natural (non-laboratory) conditions will also be important.

Continuing studies on agrochemical exposure that include evaluations of the gut microbiota as part of the studies on honey bee health, fitness, behavior, immune response, and disease susceptibility are needed as honey bee colonies continue to collapse at unprecedent rates (Sánchez-Bayo and Wyckhuys, 2019). Understanding how chemicals like zinc affect bees is essential to guide agricultural practices that effectively support ecosystem health.

## Data Availability Statement

16S rRNA sequencing data will be made publicly available at NCBI SRA upon publication. Raw survivorship data is publicly available via FigShare: <u>10.6084/m9.figshare.21692435</u>

## Acknowledgements

This study was financed in part by the Coordenação de Aperfeiçoamento de Pessoal de Nível Superior - Brasil (CAPES) - Finance Code 001, as part of the CAPES-PrInt Project “Omic sciences applied to the prevention of antimicrobial resistance at the human-animal-environment interface-a one health approach” (88881.311776/2018-01; Theme III: Caatinga Biome, Biodiversity and Sustainability), and scholarship to KOS (88887.465824/2019-00). This research was also funded by Conselho Nacional de Pesquisa e Desenvolvimento (CNPq, 3136678/2020-0) and Financiadora de Estudos e Projetos (FINEP) and OSU IDI Trainee Research Award.

## Author Contributions

Conceptualization: VLH, KOS, DR, RJ

Methodology (Sample Collection and Testing): KOS, DR, RJ, CJBO

Investigation (Analysis): KOS, DR

Visualization: KOS, DR

Funding acquisition: VLH, KOS, DR, RJ

Project administration: VLH, RJ, KOS, DR

Supervision / Consultation: VLH, RJ, AER, MVL

Writing – original draft: KOS, DR, VLH

Writing – review & editing: All

## Supplementary Figures

**Supplementary Figure 1.**
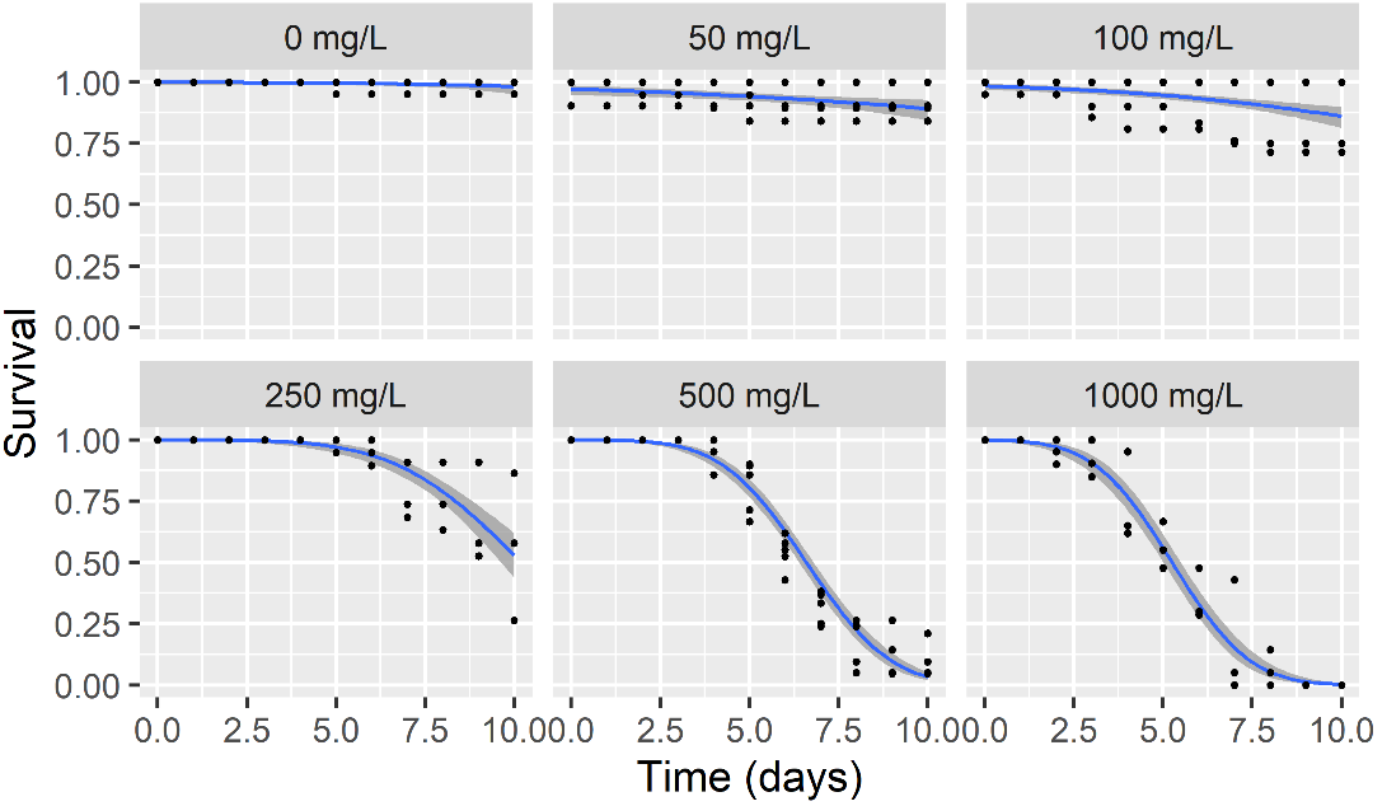
Survival rates of adult honey bees fed zinc at varying concentrations over ten days. Zinc concentrations included 0 (negative control), 50, 100, 250, 500, and 1000 mg/L. Points represent rates of honey bee survival. Lines and shaded regions represent the median survival predictions and 95% confidence regions, respectively, of probit models fitted to each dataset. Zinc exposure had a statistically significant effect on survival (likelihood ratio test, p < 2e-16, df = 22). When focusing on just the field-relevant concentrations (0, 50, and 100 mg/L), exposure still had a statistically significant effect on survival (likelihood ratio test, p = 0.007, df = 11).

**Supplementary Figure 2.**
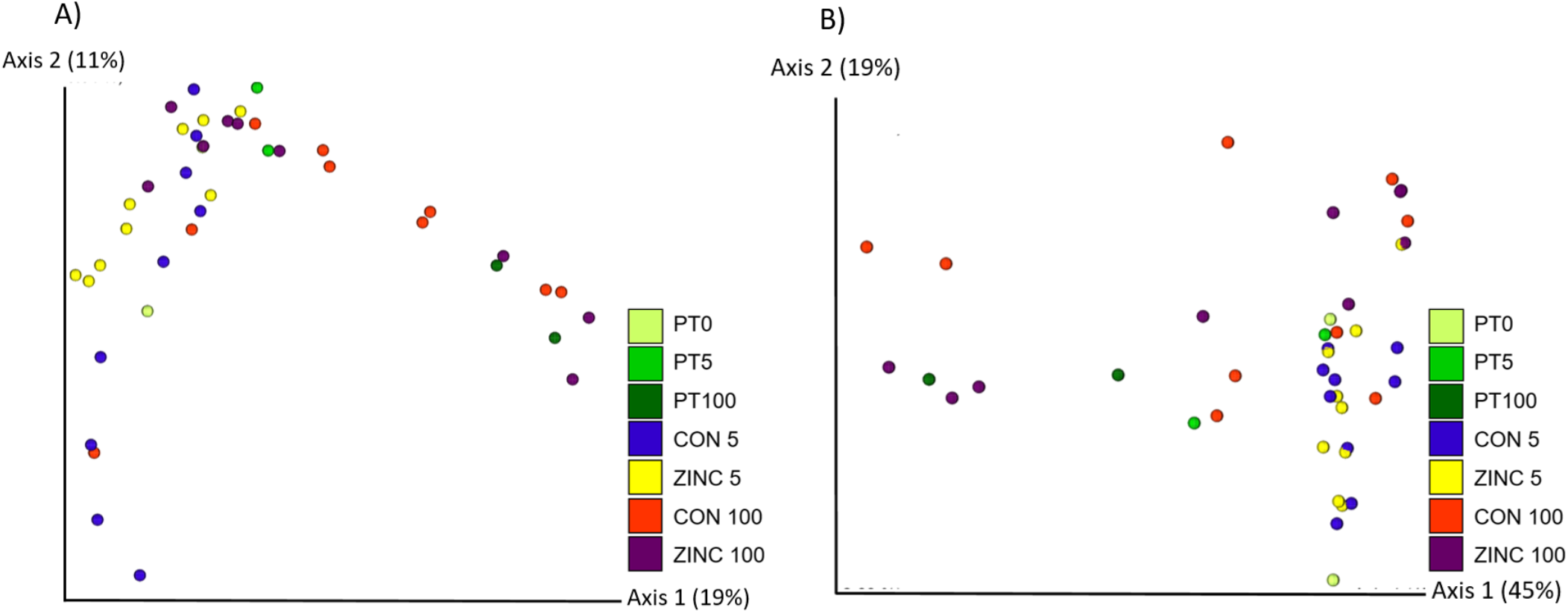
Gut microbial composition by treatment. Bee gut microbial composition differed significantly by treatment based on **A**) unweighted UniFrac (PERMANOVA: *p =* 0.001) and **B**) weighted UniFrac distance matrices (PERMANOVA: *p =* 0.006). These differences were largely driven by emerge day as opposed to zinc treatment (also see **Supplementary Tables 9,10**). PT0 = Pretreatment bees that emerged on Oct. 3; PT5 = Pretreatment bees that emerged on Oct 7^th^ – the same day as CON 5 and ZINC 5 bees; PT100 = Pretreatment bees that emerged on Oct. 11^th.^ – the same day as CON 100 and ZINC 100 bees); CON 5 = Bees that emerged on the same day as ZINC 5 bees but treated with no zinc; ZINC 5 = Bees treated with 5 mg/L zinc; CON 100 = Bees that emerged on the same day as ZINC 100 bees but treated with no zinc; ZINC 100 = Bees treated with 100 mg/L zinc.

**Supplementary Figure 3.**
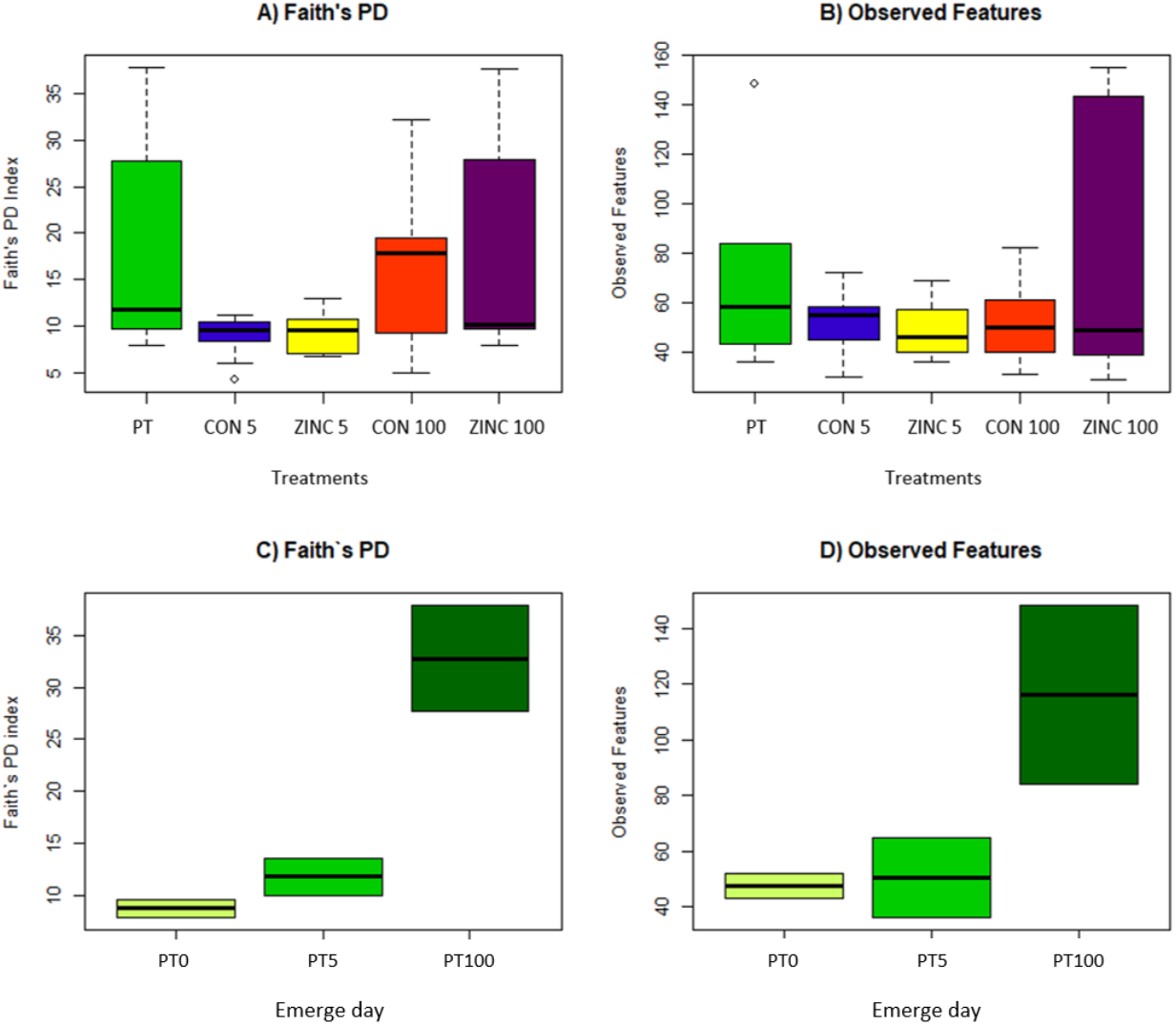
Microbial diversity by treatment. Gut microbial diversity did not differ significantly in honey bees by treatment or emerge day as measured by **A, C**) Faith’s PD (Kruskal Wallis, *p =* 0.17) and **B, D**) Observed Features (Kruskal Wallis, *p =* 0.9). An increase in microbial diversity, albeit non-significant, is observed by emerge day. Box plots show outliers, first and third quartiles (lower and upper edges), and highest, lowest and median values (horizontal black dash). PT = All Pretreatment (Day 0) bees including those that emerged on Oct. 3, 7, and 11; PT0 = Pretreatment bees that emerged on Oct. 3; PT5 = Pretreatment bees that emerged on Oct 7^th^ – the same day as CON 5 and ZINC 5 bees; PT100 = Pretreatment bees that emerged on Oct. 11^th.^ – the same day as CON 100 and ZINC 100 bees); CON 5 = Bees that emerged on the same day as ZINC 5 bees but treated with no zinc; ZINC 5 = Bees treated with 5 mg/L zinc; CON 100 = Bees that emerged on the same day as ZINC 100 bees but treated with no zinc; ZINC 100 = Bees treated with 100 mg/L zinc.

**Supplementary Figure 4.**
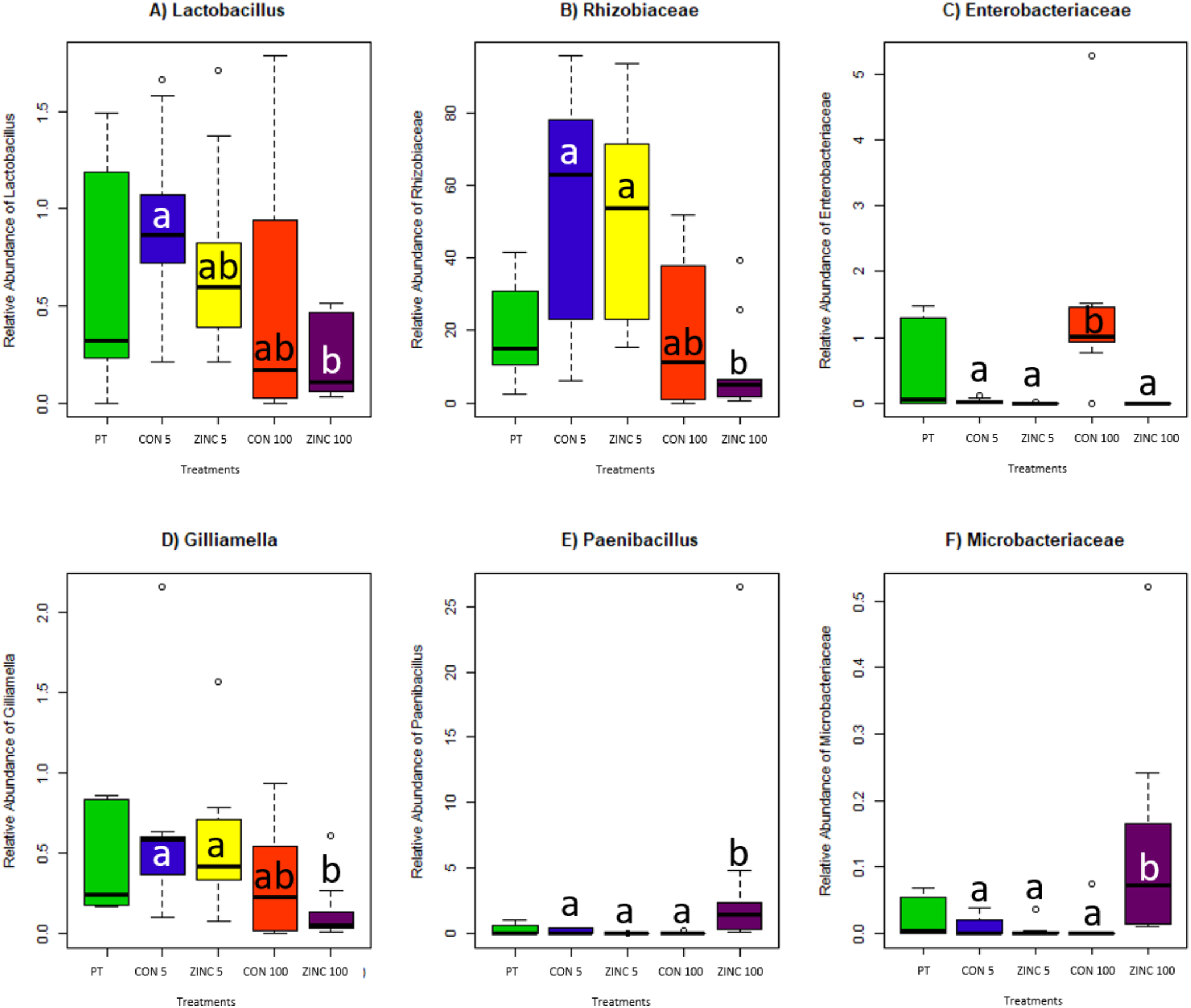
Differentially abundant microbiota by treatment. Relative abundances of microbiota that were differentially abundant (ANCOM) by treatment at the L7 (roughly species) level between groups: **A**) *Lactobacillus*, **B**) *Bifidobacterium*, **C**) Rhizobiaceae, **D**) *Gilliamella*, **E**) *Tyzzerella*, **F**) *Streptomyces*, **G**) *Paenibacillus*, **H**) Enterobacteriaceae, **I**) Proteobacteria. Box plot shows outliers, first and third quartiles (lower and upper edges), and highest, lowest and median values (horizontal black dash). Groups that share the same letter do not differ significantly (Wilcoxon rank sum test with continuity correction). PT = All Pretreatment (Day 0) bees including those that emerged on Oct. 3, 7, and 11; CON 5 = Bees that emerged on the same day as ZINC 5 bees but treated with no zinc; ZINC 5 Bees treated with 5 mg/L zinc; CON 100 = Bees that emerged on the same day as ZINC 100 bees but treated with no zinc; ZINC 100 = Bees treated with 100 mg/L zinc.

## Supplementary Tables

**Supplementary Table 1.**
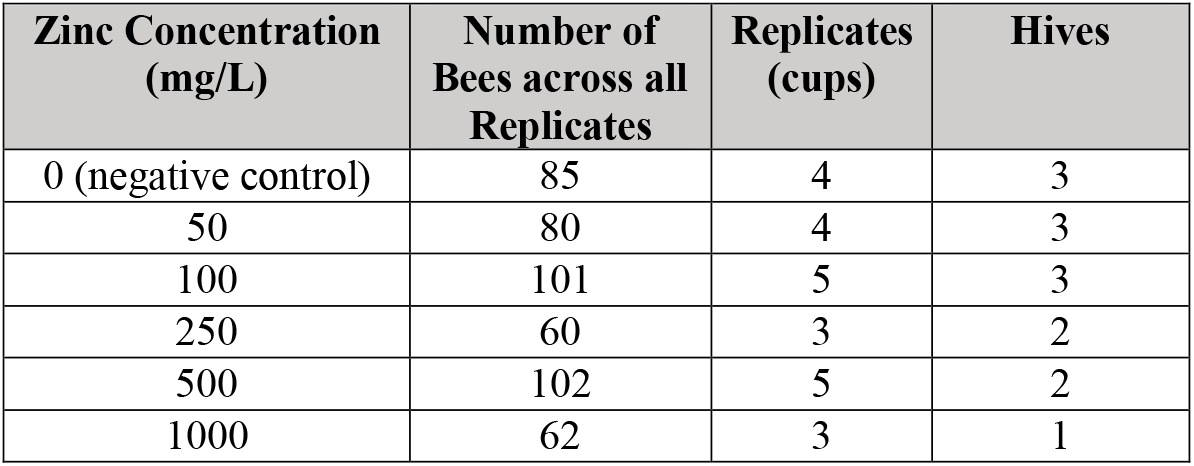
Zinc oral toxicity assays. The number of bees, replicates, and hives in each treatment group of the zinc oral toxicity trials. The final number of hives varied between 1-3 after omitting replicates with control groups that exhibited survival < 85% (OECD, 2017).

**Supplementary Table 2.**
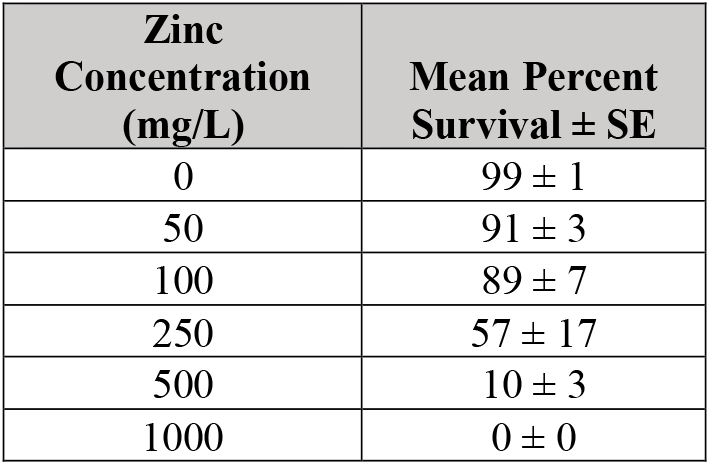
Mean rates of survival observed during oral toxicity assays after 10 days.

**Supplementary Table 3.** Honey bee survival duration estimates based on zinc exposure. LT50 is the estimated lethal effect times to reach 50% mortality. The LT50 decreased (fewer days) with increasing zinc concentrations.

| Zinc Concentration (mg/L) | LT50 Estimate (days) | Confidence Limits | Standard Error | $\chi^2$ Goodness-of-Fit Statistic | df |
| --- | --- | --- | --- | --- | --- |
| 50 | 24.89 | 18.05 - 48.13 | 5.37 | 43.92 | 42 |
| 100 | 19.39 | 14.35 - 38.7 | 2.48 | 134.31 | 53 |
| 250 | 10.20 | 9.51 - 11.29 | 0.34 | 43.87 | 31 |
| 500 | 6.59 | 6.37 - 6.82 | 0.1 | 65.01 | 53 |
| 1000 | 5.25 | 4.95 - 5.55 | 0.12 | 43.36 | 31 |

**Supplementary Table 4.** P-values from pairwise log-rank tests of differences in the overall rates of survival observed between treatment groups receiving varying zinc concentrations. All pairwise comparisons were significant (p << 0.05) except the comparison between the 50 and 100 mg/L groups.

| Zinc Concentration (mg/L) |  |  |  |  |  |
| --- | --- | --- | --- | --- | --- |
|  | 0 | 50 | 100 | 250 | 500 |
| 50 | 2.33E-09 | NA | NA | NA | NA |
| 100 | 2.23E-10 | ~1 | NA | NA | NA |
| 250 | 2.32E-21 | 0.002 | 0.003 | NA | NA |
| 500 | 3.03E-90 | 4.77E-56 | 4.61E-65 | 5.45E-31 | NA |
| 1000 | 2.99E-112 | 2.38E-74 | 1.74E-86 | 4.94E-46 | 2.75E-05 |

**Supplementary Table 5.** Total and bacterial DNA concentrations and read counts. Total and bacterial DNA concentrations and number of 16S reads in each sample. (16S read counts are reported both before and after running DADA2. PT = All Pretreatment (Day 0) bees including those that emerged on Oct. 3, 7, and 11; PT0 = Pretreatment bees that emerged on Oct. 3; PT 5 = Pretreatment bees that emerged on Oct 7^th^ – the same day as CON 5 and ZINC 5 bees; PT 100 = Pretreatment bees that emerged on Oct. 11^th.^ – the same day as CON 100 and ZINC 100 bees); CON 5 = Bees that emerged on the same day as ZINC 5 bees but treated with no zinc; ZINC 5 = Bees treated with 5 mg/L zinc; CON 100 = Bees that emerged on the same day as ZINC 100 bees but treated with no zinc; ZINC 100 = Bees treated with 100 mg/L zinc; D3 = Day 3, D6 = Day 6, D9 = Day 9. * = Excluded from analyses as variation between all replicates was greater than 3%.

| Sample ID | Group | Zinc Conc. (mg/L) | Emerge Day | Time | Cycle Threshold Values | Total DNA Conc. (ng/ul) | Bacterial DNA Conc. (ng/ul) | Percent Bacterial DNA | Number of 16S Reads (raw) | Number of 16S Reads (post DADA2) | Total Absolute Cell Counts | Retained in analyses |
| --- | --- | --- | --- | --- | --- | --- | --- | --- | --- | --- | --- | --- |
| PT1 | PT / PT0 | 0 | 10/03/20 | Pretreatment | 18.16 | 141 | 2.77 | 1.96 | 115,704 | 20,752 | 558,131 | Yes |
| PT2 | PT / PT0 | 0 | 10/03/20 | Pretreatment | 21.77 | 117 | 0.58 | 0.50 | 93,268 | 4,580 | 117,514 | Yes |
| PT3 | PT / PT5 | 0 | 10/07/20 | Pretreatment | 29.36 | 136 | 0.02 | 0.02 | 94,952 | 31,368 | 4,419 | Yes |
| PT4 | PT / PT5 | 0 | 10/07/20 | Pretreatment | 24.11 | 140 | 0.21 | 0.15 | 117,458 | 7,365 | 42,718 | Yes |
| PT5 | PT / PT100 | 0 | 10/11/20 | Pretreatment | 33.60 | 140 | 0.00 | 0.00 | 129,983 | 68,083 | 707 | Yes |
| PT6 | PT / PT100 | 0 | 10/11/20 | Pretreatment | 38.71 | 157 | 0.00 | 0.00 | 137,281 | 88,988 | 78 | Yes |
| A7 | ZINC 5 | 5 | 10/07/20 | D3 | 16.78 | 229 | 5.03 | 2.20 | 91,953 | 39,853 | 1,013,624 | Yes |
| A8 | ZINC 5 | 5 | 10/07/20 | D3 | 20.32 | 128 | 1.09 | 0.85 | 88,601 | 21,952 | 219,924 | Yes |
| A9 | ZINC 5 | 5 | 10/07/20 | D3 | 17.65 | 195 | 3.46 | 1.77 | 132,705 | 36,577 | 696,909 | Yes |
| A10 | ZINC 5 | 5 | 10/07/20 | D6 | 19.45 | 144 | 1.58 | 1.10 | 146,690 | 47,571 | 384,943 | No * |
| A11 | ZINC 5 | 5 | 10/07/20 | D6 | 18.97 | 167 | 1.95 | 1.17 | 122,443 | 29,375 | 280,759 | No * |
| A12 | ZINC<br>5 | 5 | 10/07/20 | D6 | 18.76 | 219 | 2.14 | 0.98 | 132,757 | 84,203 | 430,497 | Yes |
| A13 | ZINC<br>5 | 5 | 10/07/20 | D9 | 16.44 | 214 | 5.82 | 2.72 | 128,878 | 40,413 | 1,173,826 | Yes |
| A14 | ZINC<br>5 | 5 | 10/07/20 | D9 | 17.48 | 192 | 3.71 | 1.93 | 135,517 | 61,750 | 748,191 | Yes |
| A15 | ZINC<br>5 | 5 | 10/07/20 | D9 | 20.07 | 174 | 1.21 | 0.70 | 114,409 | 6,452 | 244,561 | Yes |
| B16 | ZINC<br>100 | 100 | 10/11/20 | D3 | 18.17 | 154 | 2.76 | 1.79 | 113,731 | 28,476 | 615,873 | No * |
| B17 | ZINC<br>100 | 100 | 10/11/20 | D3 | 35.59 | 124 | 0.00 | 0.00 | 106,334 | 56,532 | 299 | Yes |
| B18 | ZINC<br>100 | 100 | 10/11/20 | D3 | 19.54 | 155 | 1.53 | 0.99 | 137,343 | 105,651 | 308,074 | Yes |
| B19 | ZINC<br>100 | 100 | 10/11/20 | D6 | 24.66 | 166 | 0.17 | 0.10 | 77,309 | 4,544 | 33,627 | Yes |
| B20 | ZINC<br>100 | 100 | 10/11/20 | D6 | 19.54 | 177 | 1.53 | 0.86 | 67,922 | 3,848 | 307,770 | Yes |
| B21 | ZINC<br>100 | 100 | 10/11/20 | D6 | 32.86 | 162 | 0.00 | 0.00 | 120,927 | 32,924 | 975 | Yes |
| B22 | ZINC<br>100 | 100 | 10/11/20 | D9 | 37.87 | 182 | 0.00 | 0.00 | 123,715 | 69,452 | 112 | Yes |
| B23 | ZINC<br>100 | 100 | 10/11/20 | D9 | 19.08 | 176 | 1.86 | 1.06 | 126,685 | 101,403 | 375,157 | Yes |
| B24 | ZINC<br>100 | 100 | 10/11/20 | D9 | 21.25 | 183 | 0.73 | 0.40 | 135,415 | 107,692 | 146,927 | Yes |
| C25 | CON 5 | 0 | 10/07/20 | D3 | 19.84 | 144 | 1.34 | 0.93 | 105,481 | 25,491 | 269,872 | Yes |
| C26 | CON 5 | 0 | 10/07/20 | D3 | 18.92 | 180 | 1.99 | 1.11 | 97,260 | 24,495 | 401,622 | Yes |
| C27 | CON 5 | 0 | 10/07/20 | D3 | 21.08 | 136 | 0.79 | 0.58 | 98,363 | 44,490 | 158,272 | Yes |
| C28 | CON 5 | 0 | 10/07/20 | D6 | 17.92 | 176 | 3.08 | 1.75 | 129,716 | 47,728 | 620,423 | Yes |
| C29 | CON 5 | 0 | 10/07/20 | D6 | 16.23 | 179 | 6.38 | 3.56 | 112,603 | 49,089 | 1,495,437 | No * |
| C30 | CON 5 | 0 | 10/07/20 | D6 | 16.42 | 191 | 5.87 | 3.08 | 125,249 | 59,330 | 1,184,372 | Yes |
| C31 | CON 5 | 0 | 10/07/20 | D9 | 17.89 | 222 | 3.12 | 1.40 | 79,066 | 31,636 | 628,082 | Yes |
| C32 | CON 5 | 0 | 10/07/20 | D9 | 18.66 | 183 | 2.24 | 1.22 | 91,428 | 36,709 | 450,635 | Yes |
| C33 | CON 5 | 0 | 10/07/20 | D9 | 20.60 | 162 | 0.96 | 0.59 | 103,253 | 15,614 | 192,676 | No * |
| D34 | CON<br>100 | 0 | 10/11/20 | D3 | 19.92 | 167 | 1.29 | 0.77 | 119,104 | 5,379 | 260,604 | Yes |
| D35 | CON<br>100 | 0 | 10/11/20 | D3 | 19.84 | 147 | 1.34 | 0.91 | 108,771 | 7,940 | 269,888 | Yes |
| D36 | CON<br>100 | 0 | 10/11/20 | D3 | 20.72 | 195 | 0.92 | 0.47 | 121,383 | 10,367 | 184,862 | Yes |
| D37 | CON<br>100 | 0 | 10/11/20 | D6 | 20.05 | 157 | 1.22 | 0.78 | 109,498 | 6,820 | 246,472 | Yes |
| D38 | CON<br>100 | 0 | 10/11/20 | D6 | 18.35 | 255 | 2.55 | 1.00 | 104,853 | 23,899 | 515,108 | Yes |
| D39 | CON<br>100 | 0 | 10/11/20 | D6 | 19.62 | 185 | 1.47 | 0.80 | 139,057 | 10,041 | 297,290 | Yes |
| D40 | CON<br>100 | 0 | 10/11/20 | D9 | 19.52 | 140 | 1.54 | 1.10 | 131,680 | 23,263 | 310,587 | Yes |
| D41 | CON<br>100 | 0 | 10/11/20 | D9 | 18.42 | 149 | 2.47 | 1.66 | 133,753 | 43,962 | 498,253 | Yes |
| D42 | CON<br>100 | 0 | 10/11/20 | D9 | 20.77 | 167 | 0.90 | 0.54 | 148,812 | 5,324 | 180,869 | Yes |

**Supplementary Table 6.** Taxa that were identified by decontam as potential contaminants. These taxa were bioinformatically removed prior to analyses.

| Taxa ID | Description |
| --- | --- |
| 354acc009b4d5743f420af8925691527 | Romboutsia sp. CE17 chromosome, complete genome |
| 302f68ec5342caa417fa82f6bd6a6eeb | Methylobacterium sp. SO-204 gene for 16S ribosomal RNA, partial sequence |
| 05993181c81cb6e09bf67c0093fb2595 | Cutibacterium acnes strain 3265 16S ribosomal RNA gene, partial sequence |
| 9fcd9366dd83ecf153270aa8c8d4e609 | Methylorubrum extorquens strain B44 16S ribosomal RNA gene, partial sequence |
| b8ecf6ec5cc4215ab3954b72f41f0912 | Uncultured bacterium clone A1435 16S ribosomal RNA gene, partial sequence |

**Supplementary Table 7.** Evaluating effects of treatment (ZINC vs. CONTROL) and time (days) on gut microbial composition (Bray Curtis -PERMANOVA). Only treatment was significant.

|  | Df | SumsOfSqs | MeanSqs | F.Model | R2 | Pr(>F) |
| --- | --- | --- | --- | --- | --- | --- |
| **Treatment** | 4 | 3.262155 | 0.815539 | 3.718791 | 0.280026 | 0.001 |
| **Time** | 1 | 0.308681 | 0.308681 | 1.407563 | 0.026497 | 0.201 |
| **Treatment:Time** | 3 | 0.841663 | 0.280554 | 1.279305 | 0.072249 | 0.203 |
| **Residuals** | 33 | 7.236970 | 0.219302 | NaN | 0.621227 | NaN |
| **Total** | 41 | 11.649469 | NaN | NaN | 1.000000 | NaN |

**Supplementary Table 8.** Evaluating effects of treatment (ZINC vs. CONTROL) and time (days) on gut microbial composition (Jacaard -PERMANOVA). Only treatment was significant.

|  | Df | SumsOfSqs | MeanSqs | F.Model | R2 | Pr(>F) |
| --- | --- | --- | --- | --- | --- | --- |
| **Treatment** | 4 | 2.702407 | 0.675602 | 2.310244 | 0.198455 | 0.001 |
| **Time** | 1 | 0.285827 | 0.285827 | 0.977395 | 0.020990 | 0.509 |
| **Treatment:Time** | 3 | 0.978558 | 0.326186 | 1.115404 | 0.071862 | 0.146 |
| **Residuals** | 33 | 9.650435 | 0.292437 | NaN | 0.708693 | NaN |
| **Total** | 41 | 13.617227 | NaN | NaN | 1.000000 | NaN |

**Supplementary Table 9.** Bray Curtis PERMANOVA pairwise comparisons. Notably, CON 100 and ZINC 100 groups do not differ. CON 5 and ZINC 5 groups also do not differ. However, CON 100 and ZINC 100 groups differ significantly from CON 5 and ZINC 5 groups. Treatment groups included: PT = Pretreatment (Day 0), CON 5 = Bees that emerged on the same day as ZINC 5 bees but treated with no zinc, ZINC 5 = Bees treated with 5 mg/L zinc, CON 100 = Bees that emerged on the same day as ZINC 100 bees but treated with no zinc, ZINC 100 = Bees treated with 100 mg/L zinc.

| Group 1 | Group 2 | Sample size | Permutations | pseudo-F | p-value | q-value |
| --- | --- | --- | --- | --- | --- | --- |
| ZINC 100 | PT | 15 | 999 | 1.94 | 0.07 | 0.088 |
| ZINC 100 | CON 100 | 18 | 999 | 1.68 | 0.175 | 0.194 |
| ZINC 100 | CON 5 | 18 | 999 | 6.26 | 0.001 | 0.003 |
| ZINC 5 | ZINC 100 | 18 | 999 | 6.5 | 0.001 | 0.003 |
| ZINC 5 | PT | 15 | 999 | 4.15 | 0.001 | 0.003 |
| ZINC 5 | CON 100 | 18 | 999 | 3.57 | 0.006 | 0.010 |
| ZINC 5 | CON 5 | 18 | 999 | 0.79 | 0.514 | 0.514 |
| PT | CON 100 | 15 | 999 | 1.0 | 0.039 | 0.056 |
| PT | CON 5 | 15 | 999 | 5.05 | 0.001 | 0.003 |
| CON 5 | CON 100 | 18 | 999 | 4.35 | 0.004 | 0.008 |

**Supplementary Table 10.** Jaccard PERMANOVA pairwise comparisons. All groups differed significantly. PT = Pretreatment (Day 0), CON 5 = Bees that emerged on the same day as ZINC 5 bees but treated with no zinc, ZINC 5 = Bees treated with 5 mg/L zinc, CON 100 = Bees that emerged on the same day as ZINC 100 bees but treated with no zinc, ZINC 100 = Bees treated with 100 mg/L zinc.

| Group 1 | Group 2 | Sample size | Permutations | pseudo-F | p-value | q-value |
| --- | --- | --- | --- | --- | --- | --- |
| ZINC 100 | PT | 15 | 999 | 1.56 | 0.004 | 0.004 |
| ZINC 100 | CON 100 | 18 | 999 | 2.17 | 0.001 | 0.001 |
| ZINC 100 | CON 5 | 18 | 999 | 2.68 | 0.001 | 0.001 |
| ZINC 5 | ZINC 100 | 18 | 999 | 3.34 | 0.001 | 0.001 |
| ZINC 5 | PT | 15 | 999 | 2.08 | 0.001 | 0.001 |
| ZINC 5 | CON 100 | 18 | 999 | 3.21 | 0.001 | 0.001 |
| ZINC 5 | CON 5 | 18 | 999 | 1.88 | 0.001 | 0.001 |
| PT | CON 100 | 15 | 999 | 1.47 | 0.022 | 0.022 |
| PT | CON 5 | 15 | 999 | 1.89 | 0.001 | 0.001 |
| CON 5 | CON 100 | 18 | 999 | 2.48 | 0.001 | 0.001 |

**Supplementary Table 11.** Core microbiota. L7 – or roughly species level taxa - that were present in 100% of samples across all treatment groups.

| Feature ID |
| --- |
| D_1 Proteobacteria;D_2 Alphaproteobacteria;D_3 Rhizobiales;D_4 Rhizobiaceae |
| D_1 Actinobacteria;D_2 Actinobacteria;D_3 Bifidobacteriales;D_4 Bifidobacteriaceae;<br>D_5 Bifidobacterium |
| D_1 Proteobacteria;D_2 Alphaproteobacteria;D_3 Acetobacterales;D_4 Acetobacteraceae;<br>D_5 Commensalibacter |
| D_1 Firmicutes;D_2 Bacilli;D_3 Lactobacillales;D_4 Lactobacillaceae;D_5 Lactobacillus |

**Supplementary Table 12.** Differentially abundant taxa (ANCOM) at the genera (L7) level by treatment.

| Feature ID |  | W |
| --- | --- | --- |
| D_0 Bacteria;D_1 Firmicutes;D_2 Bacilli;D_3 Lactobacillales;D_4 Lactobacillaceae;D_5 Lactobacillus | Decreased in PT 100 | 409 |
| D_0 Bacteria;D_1 Proteobacteria;D_2 Alphaproteobacteria;D_3 Rhizobiales;D_4 Rhizobiaceae | Decreased in PT 100 | 409 |
| D_0 Bacteria;D_1 Proteobacteria;D_2 Gammaproteobacteria;D_3 Enterobacteriales;D_4 Enterobacteriaceae | Increased in CON 100 | 409 |
| D_0 Bacteria;D_1 Proteobacteria;D_2 Gammaproteobacteria;D_3 Orbales;D_4 Orbaceae;D_5 Gilliamella | Decreased in ZINC 100 | 408 |
| D_0 Bacteria;D_1 Firmicutes;D_2 Bacilli;D_3 Bacillales;D_4 Paenibacillaceae;D_5 Paenibacillus | Increased in ZINC 100 | 408 |
| D_0 Bacteria;D_1 Actinobacteria;D_2 Actinobacteria;D_3 Micrococcales;D_4 Microbacteriaceae | Increased in ZINC 100 | 375 |

